# Lymphatic dynamics visualized by native transient hypoxia imaging in areas of tissue damage, edema and in sentinel lymph nodes

**DOI:** 10.1101/2024.09.27.615461

**Authors:** Marien I. Ochoa, Matthew S. Reed, Weifeng Zeng, Samuel O. Poore, Tayyaba Hasan, Brian W. Pogue

## Abstract

Imaging lymphatic compartments and their function has always been challenging, yet this capability is key to understanding the dynamics of immune response and lymph dysfunction in disease states. This study reports the first ever visualization of murine lymphatic pumping and function imaged from the inherent transient hypoxia that occurs within the lymph ducts and nodes. The lymphatic system appears as one of the few naturally hypoxic areas in vivo. Hypoxia in lymphatics is detected via delayed fluorescence (DF) of endogenous protoporphyrin IX (PpIX), enabling real-time imaging. Lymph nodes and their function were localized by hypoxia transcutaneous imaging and in surgically exposed nodes, followed by correlation of localization to indocyanine green (ICG) local injection. The lymphatic pumping frequency was altered through progressive damage from mild, moderate, and severe wound injuries, and hypoxia appeared readily in the sentinel lymph nodes near tumor regions. Cyclical pumping was observed at sites of edema and in nodes near tumors. Control data from uninjured anesthetized mice showed little lymphatic contrast, whereas awake mice exhibited hypoxia localized to lymph nodes. Unlike contrast injection-based regional lymph node imaging by ICG or MRI, DF hypoxia imaging appears to provide a natural whole-body contrast mechanism, highlighting its potential for visualizing lymphatic function and associated hypoxia dynamics. The value for localization of sentinel lymph nodes and for allowing for visualization of damaged lymph has very practical potential applications.

## 1. INTRODUCTION

To date, the lymphatic network -- one of the major circulatory systems in the body alongside the cardiovascular system -- has not been fully understood.^1^ This is mainly due to its complexity and nearly invisible nature – as lymphatic channels are not easily observed by human vision in comparison to cardiovascular structures^2^. In addition, unlike the cardiovascular system, movements are passive and not coordinated through larger structures (e.g., heart pumping).^34^ The lymphatic system is a crucial part of the body’s immune response due to its role in the transport of white blood cells and providing an isolated system to protect the organism against invaders and infections. Hence understanding how the lymphatic system responds upon treatment (ex. Immunotherapy) has gained attention.^5^ Another essential function of the lymphatic system is the prevention of edema through regulation of fluids in conjunction to the cardiovascular system.^1^ Hence, the capabilities of the lymphatic system to properly regulate immune cell production and function, fat absorption, and overall fluid-balance, play a vital role in protecting the body against infections and supporting overall health. However, just as critically functional as they are, they are one of the most challenging organ systems in the body to visualize because lymph vessels are exceedingly small, and fluid that permeates them is visually clear. Studies on the active pumping mechanism of lymphatics are limited, and nearly all data is from regional injection of contrast agents to visualize the flow and pumping dynamics^6^.

At present, imaging of lymphatics has been restricted to structural imaging and localization of lymphatic structures. Hence the functionality of the lymphatic system in normal and/or disease conditions has not been fully explored. Several imaging techniques like ultrasound imaging (US)^7^, bio-impedance imaging^6^, magnetic resonance imaging^8^, lymphoscintigraphy^9^, lymphangiography^10^, dynamic contrast-enhanced MRI lymphangiography (DCMRL)^11^ and NIR/ICG lymphography^12^ have attempted to image lymphatic function. However, there is currently no established non-ionizing or minimally invasive method that can assess lymphatic function in real-time.^6^ An important and unique feature of the lymphatic system is its hypoxic nature ^13,14^, which has been seldom explored. Hypoxia in lymphatics can exist naturally, and results from lymph vessels and nodes being frequently situated in remote areas far from oxygen-carrying blood vessels. ^15,16^ Hypoxia is also expected as lymphatics are an unregulated receiving system for waste fluid and molecules. During times of high metabolic demand of oxygen in the body, it is plausible that extracellular fluid can be transiently depleted of oxygen. Previous studies have reported on the role of hypoxic conditions in the lymphatic system in promoting lymphangiogenesis and metastatic spread through the lymphatic channels.^13,15,16^

To date, imaging of oxygen content in the lymphatic network has been mainly explored in the microscopic regime for ex vivo tissues with exogenous labels.^17^ This work utilizes endogenous protoporphyrin IX (PpIX) delayed fluorescence (DF) optical imaging as method to macroscopically image hypoxia dynamics in lymphatics in vivo and in real-time. PpIX production can be enhanced by its precursor 5-ALA, which is already FDA approved for human use.^18^ Initially, the localization of lymphatics through PpIX DF optical imaging was correlated to lymphatic vessels and nodes labelled through ICG^19,20^ for uninjured mice. Lymphatic function was subsequently evaluated for mice in vivo through induced moderate and severe wound injuries, with uninjured mice serving as controls. Real time PpIX DF imaging was also performed when mice were moving. Finally, PpIX DF imaging was used to dynamically (i.e., in real-time) image the lymphatic response to pancreatic adenocarcinoma (AsPC1) tumors in vivo.

## II. RESULTS

### A. PPIX DF HYPOXIA DYNAMICS IN RELATION TO ICG AND VITAL SIGNS

To confirm that the detected DF hypoxia signals corresponded to lymphatics, simultaneous ICG lymph node imaging was used to visualize the location of the same lymph nodes closer to the wound area, as described in the Methods section. Figure 1 displays these results, where Figure 1 (a) shows the PF signal of PpIX representative of PpIX distribution and concentration on the intact mouse. It was observed that PpIX prompt fluorescence (PF) was higher at the wound site with fluorescence also present across the mouse skin. ICG injected in the right foot pad localized both popliteal (PLN) and sciatic (SLN) lymph nodes as shown in Figure 1 (b). These nodes also displayed draining activity as shown in Figure 1 (c) where frames were selected from the real-time sequence and for display purposes at 1, 2, and 3 minutes after imaging started. Figure 1(d) displays minute 4 of the real-time imaging sequence. In this sequence, the nodes that correlated to ICG are indicated with red arrows and nodes that were not located by ICG but only by PpIX DF imaging are indicated in a different color. Notably, DF PpIX imaging displayed more areas of activity across the mouse than ICG imaging. ICG injection on the footpad only reached the SLN and PLN nodes as previously reported. ^20–22^ Pulsation effects were not observed in the PLN or SLN node through ICG but were observed through PpIX DF Imaging. Quantification of these pulsations are displayed in Figure 1(e) for the group of mice. An example of the pulsation sequence of PpIX DF intensity from the PLN region is shown in Figure 1(e), in comparison to measurements acquired with a vital sign monitor, where heart rate and respiration signals were recorded over the measurement time. Lymphatic signals recovered through DF imaging of PpIX pulsed at a slower rate compared to respiration and heart rate signals at an average of 10 pulses per minute (bpm) as quantified in Figure 1(e).

**FIGURE 1.**
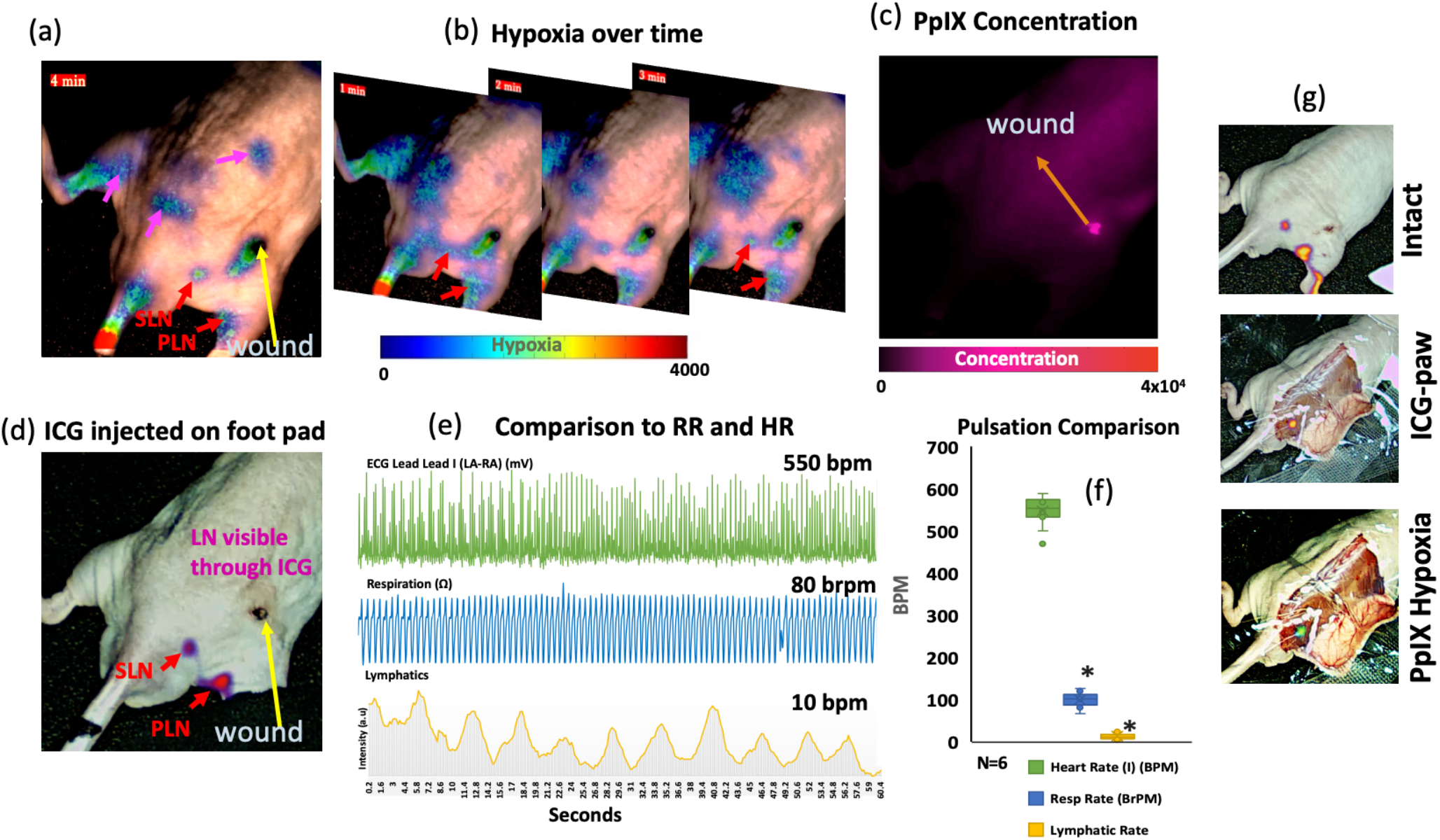
Example lymph response as recorded by PpIX DF imaging for a mouse with a moderate level wound injury. (a) PpIX prompt fluorescence (PF) channel representative of PpIX concentration. (b) ICG injected in the footpad to track neighboring nodes. (c) PpIX delayed fluorescence (DF), hypoxia channel with example frames over time. (d) PpIX DF hypoxia frame at 4 mins with lymph nodes that correlate to ICG (red arrows) and additional lymph nodes identified through PpIX DF imaging (pink arrows). (e) Quantification of beats/pulsations per minute (BPM) of PLN and SLN nodes in comparison to BPM recorded for respiration and heart rate. (f) Example of a recorded signal for heart rate, respiration rate and PLN lymph node. Real-time kinetics and movements can be appreciated in videos 1(a)-(c). Video 1(c) which represents PpIX PF displays no localization to lymphatics in comparison to PpIX DF and hypoxia in Video 1(a).

### B. MOVEMENT ENHANCES RESPONSE AND VISUALIZATION OF LYMPH NODES

Due to the passive movements expected from the lymphatic structures^3,4^, and their predicted faster function under movement, real-time PpIX DF imaging was performed on uninjured mice in movement as described in the Methods section. Results are shown in Figure 2 for a mouse under anesthesia and subsequently awake and ambulating. Figure 2(a) shows PpIX prompt fluorescence (PF) that demonstrated PpIX fluorescence throughout the mouse’s body. When the mouse ambulates (Figure 2(b) and Figure 2(d)), signals originating from lymph nodes in areas that are close to the tail and flank of the hind-legs were observed in the hypoxia channel. Signal intensity was higher than that observed when mice were under anesthesia. This was quantified for the group by averaging the overall intensity counts on the flanks of the hind-legs of the mouse over a span of 3 minutes. Quantification is shown in Figure 2(e) where PpIX DF intensity counts were higher at the flank and hind-leg regions when the mouse was in movement. Of note, Figure 2(c) showed that when the intensity is lower due to anesthetic effects, dynamic signals were also captured from the chest region. The base of the tail displayed hypoxia signals for both anesthetized and non-anesthetized mice.

**FIGURE 2.**
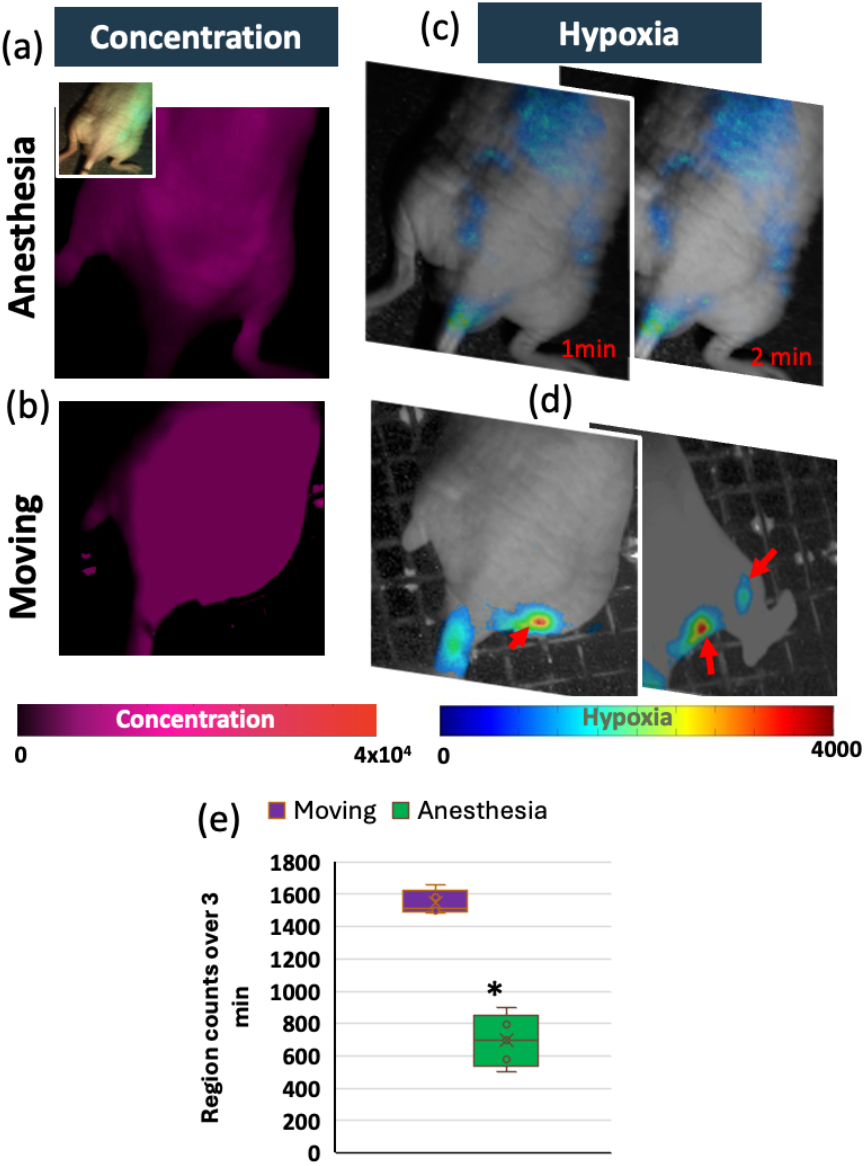
Example of lymphatic response as recorded with PpIX DF imaging for a non-injured mouse with and without anesthesia (awake and moving). (a) PpIX prompt fluorescence (PF) channel representative of PpIX concentration. White light image provided. (b) PF for an example frame at 2 minutes of the mouse without anesthesia. (c) PpIX delayed fluorescence (DF), hypoxia channel with example frames over time. (d) PpIX DF hypoxia for moving mouse with red arrows pointing at identified lymph nodes. (e) Quantification of intensity counts recorded at hind leg and back regions of the mouse over a period of 3 minutes (N = 4). Hypoxia images have been overlayed over the gray-scale white-light image for visualization purposes. Real-time kinetics and movements can be appreciated in videos 2(a)-(b).

### C. LYMPHATIC RESPONSE IN THE PRESENCE OF VISIBLE SWELLING

The experiments described in Figures 1 and 2 explored the behavior of DF PpIX signals from lymphatics in non-injury conditions and in a moderate wound injury model – where minimal visible inflammation was observed. The lymphatic behavior in areas of visible swelling was also studied in a mouse model where a more severe injury was inflicted, as described in Methods and Materials and summarized in Figure 3. Results for mouse models (N=4) of visible swelling are summarized in Figure 3. Figure 3(a) demonstrates an example mouse model of major wound injury, where the periphery of the wound was visibly inflamed. Figure 3(b) shows the PF signal of PpIX, where the recorded signal intensity (and correspondingly, measured PpIX concentration) was higher at the outside and lateral area of the wound, covering a larger area than the visibly inflamed region shown on Figure 3(a). In this example, vials of oxygenated and deoxygenated PpIX are shown within the image frame as control data. Both normal and deoxygenated vials possessed a PF component as shown in Figure 3(b) and as previously reported for in vitro samples. ^23^ PF frames at 1, 4, 9 and 13 minutes after the acquisition start are shown in Figure 3(b). Figure 3(c) displays example images of the PpIX DF real-time sequence. Frames at 1, 2, 4, 7, 9, 12 and 13 minutes are shown to illustrate the recorded intensity fluctuations. For this example, PLN and SLN pulsations were observed on both the right and left legs of the mouse. Of note, observed pulsations did not occur at the same time for each node as indicated by the red arrows in the frames displayed in Figure 3. Besides observed nodes, which are highlighted with red arrows, pulsations were observed in a larger area located at the left side of the wound, close to the visible swelling region. The intensity fluctuation of this area over time is depicted in Figure 3 (d). A total of 7 pulsations were quantified for a period of 60 seconds or a respective frequency of *∼* 0.1 Hz. Quantification of the frequency of pulsation across different time-points after 5-ALA injection for selected area and area of PLN nodes is provided in Figure 4.

**FIGURE 3.**
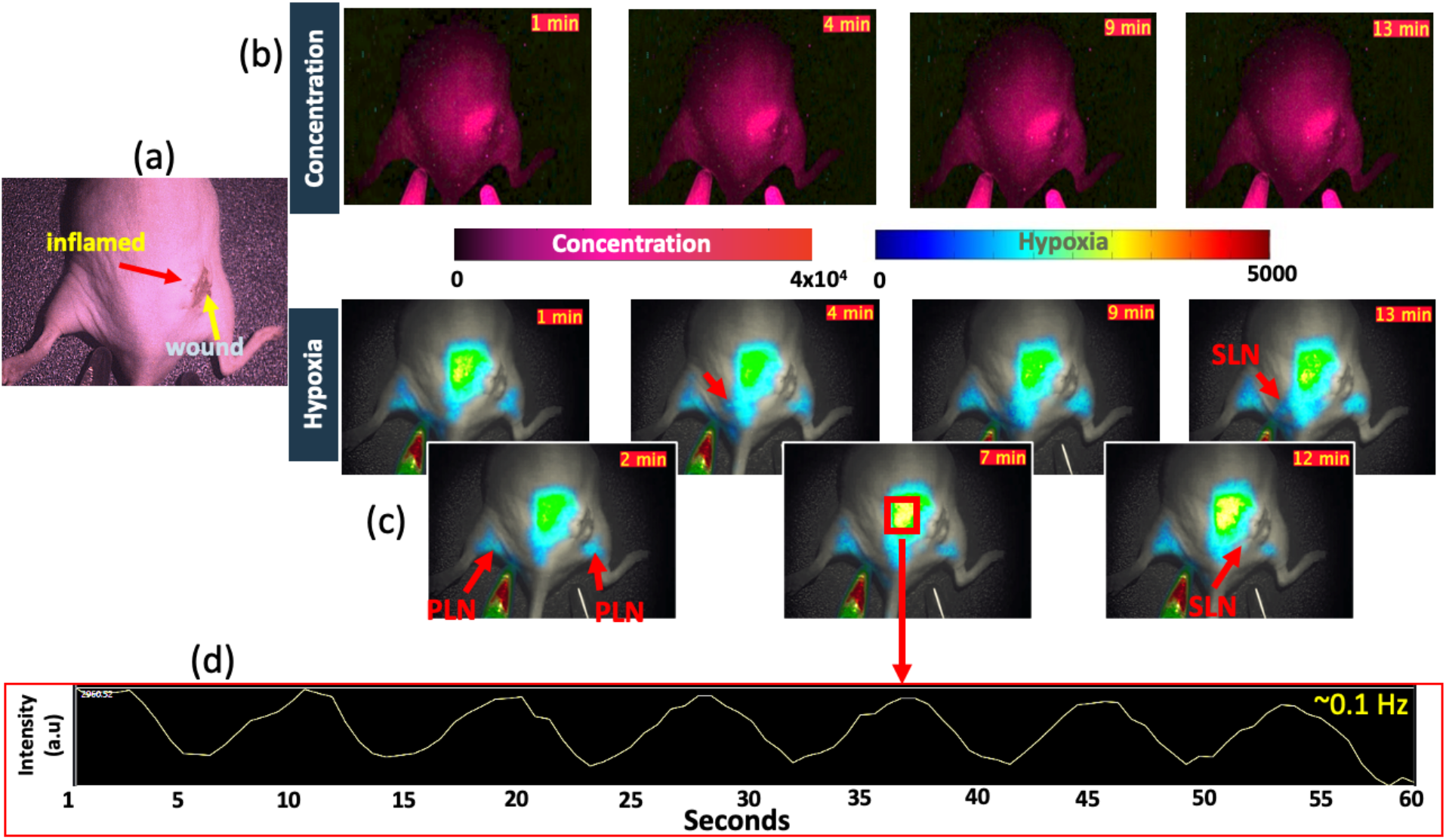
Example of lymphatic response as recorded with PpIX DF imaging for a severe injured mouse model. (a) PpIX prompt fluorescence (PF) channel representative of PpIX concentration. Whitelight image provided. (b) PF for an example frame at 2 minutes of the mouse without anesthesia. (c) PpIX delayed fluorescence (DF), hypoxia channel with example frames over time. (d) PpIX DF hypoxia for moving mouse with red arrows pointing at identified lymph nodes. (e) Quantification of intensity counts recorded at hind leg and back regions of the mouse over a period of 3 minutes. Real-time kinetics and movements can be appreciated in videos 3(a)-(b).

**FIGURE 4.**
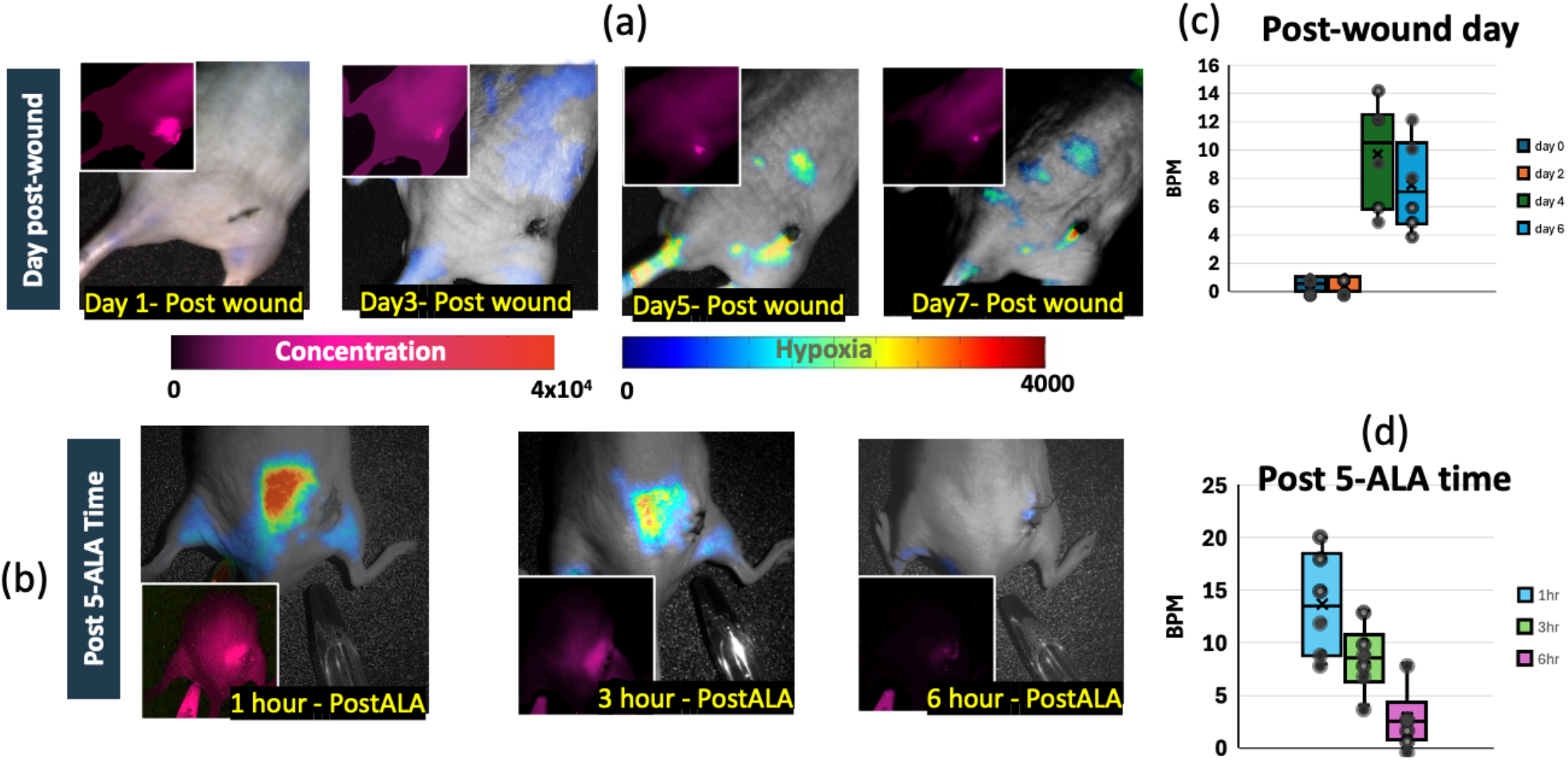
Example of lymphatic response seen in DF Hypoxia for both moderate and severe injured mouse model. (a) Hypoxia channel overlayed on white light image for a mouse injected and imaged every two days. PpIX prompt fluorescence (PF) channel representative of PpIX concentration is shown on the upper left corner for comparison. (b) Hypoxia channel overlayed on white light image of a mouse with inflammation, imaged at 1, 3, and 6 hours post 5-ALA injection. PF image displayed in left bottom corner for comparison. (c) Quantification of recorded pulses for the group (N=6) in beats per minute (BPM) and quantified in the SLN and PLN regions. (d) Quantification of recorded pulses for the group (N=4) in beats per minute (BPM) and quantified in the SLN and PLN regions for post 5-ALA injection time-points. Real-time kinetics and movements can be appreciated in video 4(a)-(d) for PF of days post-wound, video 4(e)-(h) for DF of days post-wound and videos 4(a)-(f) for PF and DF channels for post 5-ALA injection time.

### D. UNDERSTANDING WOUND PPIX SIGNALS OVER TIME

To obtain insight into which time-points better highlight areas of lymphatic pumping on the SLN and PLN areas, different time-points post-5ALA administration were compared for wound models of moderate and major injury. Results are summarized in Figure 4. Figure 4 (a) exemplifies both prompt and DF hypoxia signals, where the 5-ALA was injected in separate days at 1-hour post-injection. PF is representative of PpIX concentration. Results showed higher PpIX PF localization to the wound area at days 4 and 6. The DF hypoxia channel, as shown in Figure 4(a) and videos 4 (c) and (d), resulted in pulsations that were noticeable at days 4 and 5 post-injury but not on day 1 and 2. Besides quantifying pulsations, pulsating node positions correlated to PLN and/or SLN locations. Figure 4 (a) shows an example DF static image of hypoxic regions with higher intensity for days 5 and 7 with the dynamics being fully appreciated in videos 4 (a) and (b). The quantification of the recorded pulse number in beats per minute (BPM) for a group of mice with N=4, is shown in Figure 4 (c). Pulsations at days 5 and 7 went from 4 to 12 BPM in comparison to no pulsations observed on days 1 and day 2. Pulsations at SLN and PNL regions were only observed in the DF hypoxia channel and not in the PF channel. However, the PF channel displayed signals that were positioned around the wound on day 1 and localized to the wound region on day 6. Besides examining which day post-wound the pulsation signals started to be observed, the post 5-ALA administration time was also investigated. Results for mice imaged at 1, 3, and 6 hours post 5-ALA injection are displayed in Figure 4 (b). Inflamed areas and regions nearby displayed pockets of DF hypoxia signal that were also visible in the PF channel. However, the SLN and PLN showed pulsations over time that were only visible in the DF hypoxia channel. This can be appreciated in video 4 (c) for PpIX DF hypoxia. These pulsations were not observed in the PF channel as shown in Figure 4 (b) bottom left corner and video 4 (d). The signal accumulated near areas of inflammation and displayed higher intensity and pulsation rate (BPM) at the 1 hour post-injection time-point. The intensity in areas of interest (nearby inflammation, SLN and PLN areas) for the PF and DF channels decreased from 1 to 6 hours post 5-ALA injection. Pulsations (BMP) quantified in the DF channel also decreased from 1 to 6 hours post 5-ALA injection. These are dynamically displayed in videos 4 (c) and (d). Quantified pulsation frequencies for SLN and PLN regions are shown in Figure 4 (d) in terms of the beats per minute (BPM).

### E. LYMPHATIC RESPONSE IN THE PRESENCE OF A ASPC1 PANCREATIC TUMOR MODEL

The response of wound models in moderate and major injuries were explored in the previously described experiments for different post wound time-points and post 5-ALA injection times. To evaluate the behavior of DF PpIX signals from lymphatics in a disease (non-wound) state, DF functional imaging was studied in mice with AsPC1 (Pancreatic Adenocarcinoma) tumor xenografts. Results are displayed in Figure 5 (a) for imaging timepoints pre and post 5-ALA injection as well as 1, 3, 6, 9, 12, 16, 24 and 26 hours post 5-ALA injection. Results described in Figure 5 (a) display both PF (PF in upper left corner) and DF hypoxia channels. DF hypoxia images have been overlayed over whitelight images for visualization purposes. The pulsation rate in BPM was quantified and results plotted in Figure 5 (b). The dynamics can be fully appreciated in videos 5 (a-d) for dynamics at 1, 3, 9, and 12 hours post-5ALA injection in both DF and PF channels. Quantified BPM at SLN and PLN regions were higher at the 1-to-6-hour post-5ALA injection timepoints with higher BPM mean at the 3 hours post 5-ALA injection time-point.

**FIGURE 5.**
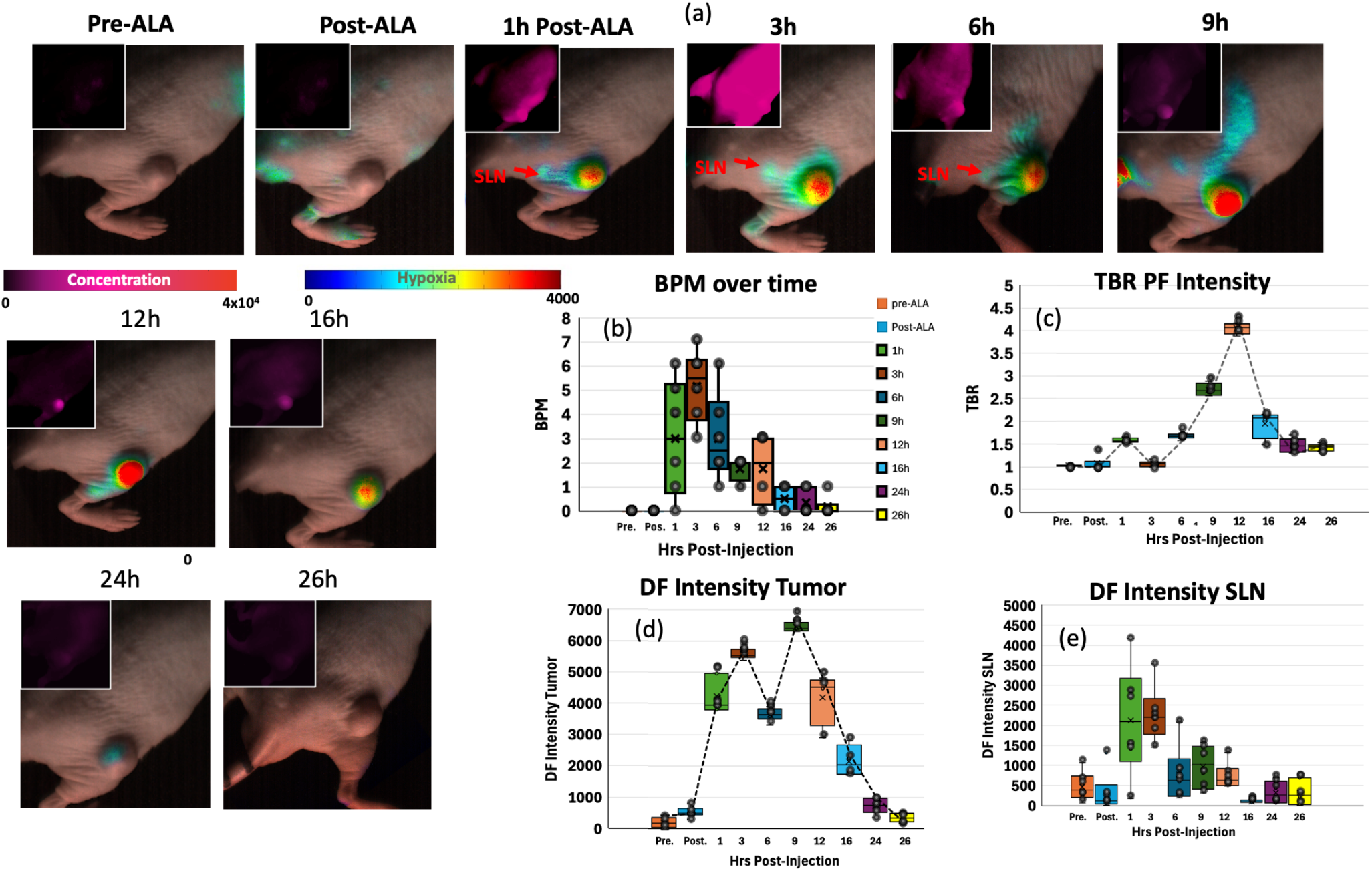
Example of lymphatic response as recorded with PpIX PF and DF imaging for a mouse with AsPC1 tumor xenograft. (a) Example sequence for time-points from before (pre-ALA), right after (post-ALA) and 1, 3, 6, 9, 12, 16, 24, and 26 hours post-5ALA injection. The Hypoxia DF channel was overlayed over the whitelight image for visualization purposes. PpIX prompt fluorescence (PF) channel representative of PpIX concentration is shown on the upper left corner for comparison. (b) Quantification of recorded pulses for the group (N=6) in beats per minute (BPM) and quantified in the SLN and PLN regions across the different time-points.(c) Quantification of tumor to background ratio (TBR) per each timepoint for prompt fluorescence (PF). (d) DF intensity on the tumor area per each time-point. (e) Quantification of DF intensities in the SLN region per each time-point. Real-time kinetics and movements can be appreciated for time-points at 1, 3, 9, 12, and 24 hours post 5-ALA in videos 5(a)-(e) for PF and videos 5(f)-(j) for DF hypoxia channel.

Quantification of tumor to background ratio (TBR) of PF is displayed in Figure 5 (c), where the TBR values peak at 12h post 5-ALA injection. DF intensity values for the tumor region were also quantified and are shown in Figure 5 (d). Results displayed higher DF intensity on the tumor region at the 9 hours post-injection time-point, with DF intensity decreasing on the subsequent time-points.

The DF intensity for the SLN area was also quantified and is displayed in Figure 5 (e). The results show a difference between the time-point where highest BPM are observed (3 hours), the time-points where highest TBR is quantified for PF (12 hours) and highest intensity quantified for the DF in the tumor region (9 hours). The time-point at which DF intensity is maximum for the SLN region (3 hours) correlates to the time-point with the highest recorded BPM.

## III. DISCUSSION

The experiments described in this manuscript explored the use of PpIX based DF hypoxia imaging as a new imaging modality to assess lymphatic function in real-time. The goal was to understand the dynamics of lymphatic function following wound injuries as well as AsPC1 tumor xenografts in mice. To our knowledge, this is the first time that hypoxia has been imaged in the lymphatic network and used as a parameter to describe lymphatic function. Results demonstrated in Figure 1 describe how hypoxia, upon wound injury, was detected in lymphatic channels and how the location of hind-leg lymph nodes were correlated to ICG based lymphatic imaging. DF lymphatic imaging correlated to SLN and PLN locations. However, in all cases, ICG only highlighted the two nodes closer to the right hind-leg as the probe was injected into the right footpad. This is one of the drawbacks of ICG based lymphatic imaging, where to visualize the full network, multiple injection sites must be targeted and will often require prior knowledge of where lymphatic channels are. ^6^ Hence, in this study ICG validation is limited to the right footpad and SLN/PLN nodes as shown in Figure 1(d). SLN and PLN nodes are targeted as wounds have been strategically positioned in the right hind leg of the mouse. Previous studies have reported on the use of SLN and PLN nodes to drain this area in both rat ^24^ and mouse models^22^. The localization of DF imaging to the SLN node was also verified by opening the skin of the mouse and comparing the location of SLN node in the DF hypoxia channel with respect to ICG. No pulsations were observed on the ICG recordings in comparison to DF lymphatic imaging, where pulsation dynamics for SLN and PLN regions are observed as seen in video 1 (a) for DF imaging and video 1(b) for ICG. Furthermore, DF imaging displayed additional features with pulsation effects as highlighted in Figure 1(a). This is likely due to systemic 5-ALA administration which distributes the probe globally instead of a localized injection site which is the case for ICG.

One key component of our study was to understand the role of respiration rate and heart rate in the observed pulsations. Observed pulsations for lymphatic regions averaged at 10 BPM rate. This value is *∼* 10 times slower than the recorded respiration rates and *∼* 55 times slower than recorded heart rates as shown in Figure 1 (e) and (f). These lymphatic BPM correlated to previously reported values where lymphatics have been observed to pump at rates from 2-15 BPM for mouse experiments^25,26^. Intensity in the lymph node areas was observed to increase with movement as shown in Figure 2, where for mice with no wound injury and under anesthesia low intensities were recorded on the SLN/PLN regions, but when the same mouse is removed from anesthesia and imaged while ambulatory, intensity values in these regions increased. In this case, only DF hypoxia intensity was quantified for 5 frames where the mouse kept a similar position – however, quantification of BPM upon movement becomes more challenging. Hence, BPM quantifications in this study were limited to when mice were under anesthesia.

When severe injuries were imaged, areas of visible inflammation as well as their proximity displayed high DF hypoxia intensity as exemplified in Figure 3(b) as well as pulsation rates at *∼* 15 BPM as shown in Figure 3(d) and Figure 3(b). Previous studies^1^ have reported on the importance of the lymphatic system in controlling inflammation by managing the drainage of extracellular fluids, inflammatory signals, and white blood cells. During inflammation, lymphatic vessels have been reported to become significantly enlarged and more permeable.^27,28^ The expansion of the vascular network has been associated with enhanced lymphatic clearance function.^29^ This phenomenon could explain why beats per minute are faster in the case of visual inflammation – however, future studies will be aimed at verifying this hypothesis. Another important aspect is the period after wound where more lymphatic pumping was observed. This was investigated and results displayed in Figure 5. Pulsations were observed at the SLN and PLN regions until Day 4 and Day 6 post-wound. This indicates that lymphatic response was not immediate after wound formation.

Previous studies have reported on increased dermal lymph vessels 3 days post-wound^30,31^, as well as lymphatic dilation observed at 1-week post-wound.^32^ Additional studies have also observed a roughly twofold increase in the number of macrophages in wounds from VEGF-A transgenic mice at Day 7 post-injury.^33^ These studies support our in vivo results observed at days 4 and 7 post-injury. However, since this work is limited to the real-time imaging aspect of lymphatic function, future work should contemplate the correlation between ex vivo analysis techniques and real-time DF imaging of lymphatics. The use of 1 hour post 5-ALA injection suggests that, even though the technique can visualize lymphatic function in real-time, the time-points after 5-ALA administration affect the output DF intensity for lymphatic regions. PpIX is eventually synthesized into heme and previous work has reported on the clearance of the probe for both delayed and PF with a respective decrease in intensity. ^34^ As shown in Figure 5, DF intensity on the lymphatic’s SLN and PLN regions is higher at 1 and 3 hours post-5ALA injection for both wounds and AsPC1 tumors. This time-point also corresponded to when DF intensity was highest at the SLN region for tumors. Since the highest TBR contrast for PF and DF in the tumor is observed past 9 hours, this gives insight that PpIX cleared by the lymphatic network is not only being cleared from the tumor area but also from other regions of the mouse. Hence, future work should look into how different 5-ALA delivery methods could enhance the signal intensity observed at the 1 to 3 hour window in lymph nodes and other lymphatic structures.

Indeed, as depicted in Figure 6, the work presented here suggests that when 5-ALA is injected promoting PpIX production, the PpIX that is produced across tissues accumulates in areas of inflammation in wounds and in the tumor. This was validated by the increase in PF intensity specific to the wound and tumor regions. It was also observed that PpIX is cleared through the lymphatics network highlighting nodes that are actively pulsing to drain the wound or tumor areas. These nodes cannot be observed through PF alone, but can be seen through the DF hypoxia channel, which is representative of hypoxia, validating the notion that lymphatics are hypoxic. This hypoxia condition has been reported as an important factor for lymphangiogenesis and metastatic progression through lymph channels^16,35^, however this is the first technique to exploit hypoxia for real-time imaging of lymphatic function.

**FIGURE 6.**
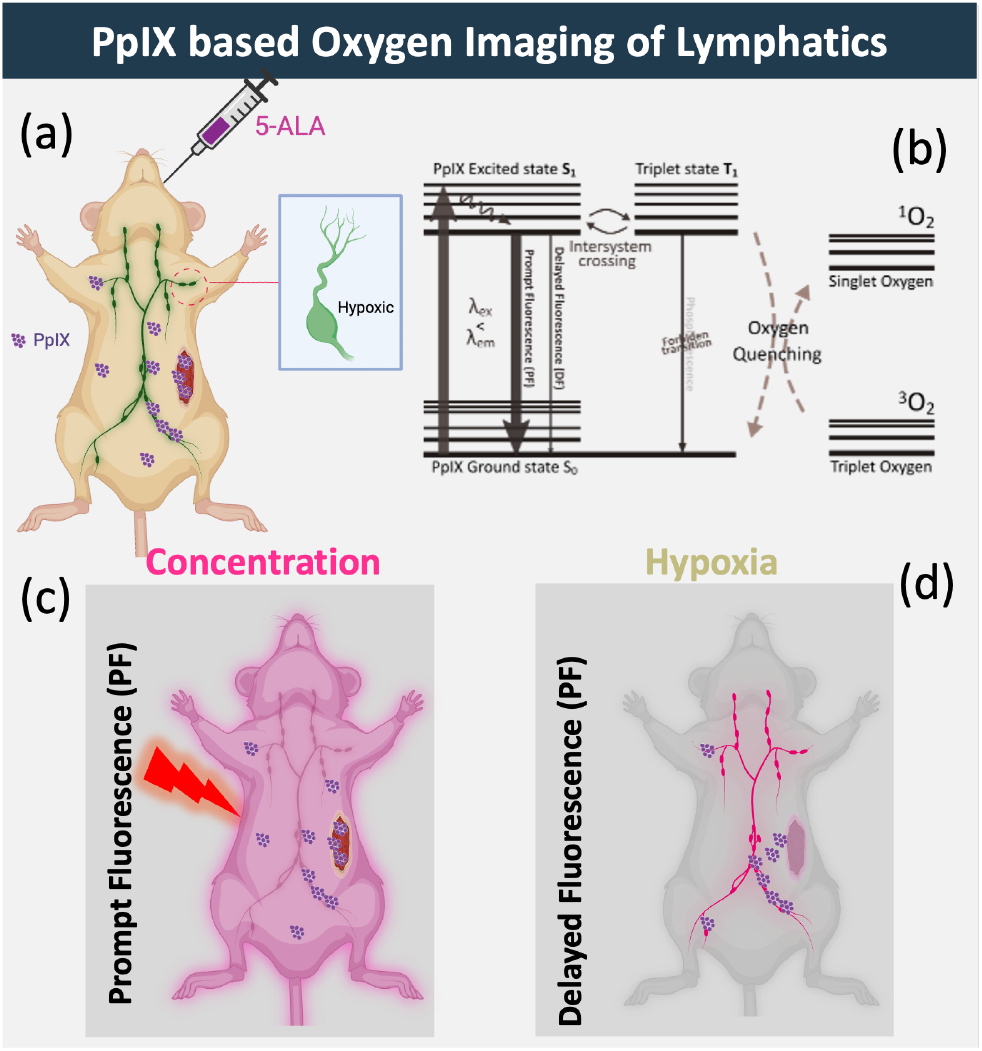
Schematic of imaging of lymphatic function through delayed fluorescence (DF) of PpIX. (a) schematic of 5-ALA retro-orbital injection. (b) Jablonski diagram displaying how delayed fluorescence is representative of hypoxia due to oxygen quenching in the triplet state. (c) Depiction of PpIX prompt fluorescence not selective of hypoxic region but representative of global PpIX concentration. (d) Delayed fluorescence component isolated through time-gated cameras and selective to the hypoxia present in the lymphatic network with PpIX as the hypoxia contrast.

One of the limitations of this study is the lack of a technique to directly compare lymphatic function in real-time and in vivo at similar frame rates as the proposed DF imaging approach. ICG was used to localize lymphatic structures connected to the footpad areas as there are extensive studies localizing hind-foot SLN and PLN connected nodes. Hence quantification of lymphatic activity is limited to these areas where we could obtain ICG validation even if DF imaging displayed additional areas of activity. While ideally we might quantify the number of nodes that become active across the whole mouse, this would require further validation across modalities with previous knowledge of the location of nodes. (ex. ultrasound, ICG imaging).

An additional observatoin was that when covered or pressed ^34^, skin also exhibits a hypoxic response (Supplementary Figure 2) as skin also has high PpIX content at specific timepoints^34,36^. This technique may help the understanding of the relationship between skin and lymphatic response of superficial nodes. The wound and tumor models in this study were positioned in the right hind-leg of mice, to control areas of drainage to the PLN and SLN nodes. It would be ideal to understand the location of active draining lymph node and dynamics across the mouse body with tumor location. To keep lymphatic structures intact, this study was limited to nodes like the PLNs and SLNs located under the skin. Even though DF was measured on some exposed nodes, one challenge is to guarantee the integrity of lymphatic channels and nodes when opening the skin, specifically for mouse models. In future studies, the use of larger animal models might better represent the nodes and lymphatic function in humans, as well as help with validation of imaging when skin is intact and during lymphatic surgery. The distribution of 5-ALA in the lymphatic channels is may be altered by different 5-ALA delivery routes (ex. oral 5-ALA formulation), although this was not tested here.

## IV. CONCLUSION

PpIX based DF hypoxia imaging of lymphatics has been presented as a fundamentally new method to visualize the lack of oxygen in lymph nodes and lymphatics vessels. To our knowledge hypoxia has not been characterized and/or used as a contrast for functional imaging of lymphatics, despite lymphatic vessels being one of the naturally hypoxic systems in the body.^13,16^ Intracellular-produced PpIX (5-ALA induced) appears to be absorbed and cleared by the lymphatic network, and exhibits a DF hypoxia component representative of the low oxygen conditions in the lymphatic fluid, especially when stressed by injury or tumor growth. Hypoxia imaging can be used to selectively visualize lymphatic structures that are being used to naturally clear PpIX. The hypoxia signal is not necessarily present in anesthetized mice, presumably due to a lack of lymph flow, but when the animal is awake and moving, the signal can be seen in the lymph nodes of the hind legs. At appropriate timepoints of 1-3hrs post 5-ALA administration, this technology can also visualize other hypoxic regions like tumors, or areas of inflammation in wounds. The time at which DF signals are higher in tumor areas is different from times at which signals are higher in lymphatics. This highlights the potential to discriminate hypoxic signals based on time-points using the same technique. Hence, this approach has the potential to uncover relationships between structures like wounds and tumors and lymphatic clearance rates. Real-time imaging enabled in vivo visualization of lymphatic flow and clearance of areas of inflammation and quantification of BPM as a metric to represent clearance rates. This is done through the use of 5-ALA, an already FDA approved contrast agent. Hence it is expected that this imaging modality will allow for lymphatic system diseases like lymphedema to be assessed from an oxygen content perspective, rather than an injection of ICG. Furthermore, will allow for the general evaluation of lymphatic activity and function upon treatments.

## V. MATERIALS AND METHODS

### A. MECHANISM OF PPIX DELAYED FLUORESCENCE IMAGING

The principle behind PpIX DF hypoxia imaging is explained in Figure 6(b). PpIX is naturally produced by organisms in the mitochondria^37^. PpIX synthesis is boosted by the administration of FDA-approved 5-ALA into the organism. ^18^ 5-ALA is a precursor in the heme pathway. Tissues containing PpIX when excited by light (350 to 650nm with 635nm use in this work), emit fluorescence from 600 to 750nm. This fluorescence can be classified into Prompt Fluorescence (PF) and Delayed Fluorescence (DF). PF of PpIX, is the most commonly used^37–40^, as its signal originates from the excited singlet state (S1) returning to the ground state. (Figure 6) PF is straightforward to excite through continuous wave illumination and can be collected through non-complex cameras (ex. cellphone cameras).^41^ PF has been effectively applied in procedures such as guided neurosurgery of glioma tumors, bacterial assesment and PDT studies^39,42,43^. On the other hand, DF of PpIX arises from emissions of S1 excited state that have replenished through reverse intersystem crossing from the triplet state (T1) (Figure 6 (b)) where oxygen quenching takes place ^44–46^. Therefore, DF fluorescence appears only in the absence of oxygen or hypoxia, thereby marking regions with low oxygen content^23,46,47^ (e.g., lymphatic systems (LS), tumors, areas of inflammation) selectively visible. This is not true for normoxic tissue, where the PpIX DF signal is quickly quenched by tissue oxygen^46^. DF has a long fluorescence lifetime decay (3ms) compared to PF’s nanosecond lifetimes (<12ns)^48,49^. Hence the DF signal, can be isolated using time-gated cameras and pulsed illumination^34^. The use of a detector grid (ex. CCD) allows for spatial mapping of delayed fluorescence (DF) representative of hypoxia. DF of PpIX has been recently explored as a method for real-time surgical guidance of tumors^34^, evaluation of oxygen content in burns^50^, and PDT assessment^46^.

### B. PPIX DELAYED FLUORESCENCE IMAGING INSTRUMENTATION

The pre-clinical DF hypoxia imaging setup is depicted in Supplementary Figure 1. Macroscopic imaging of PpIX DF hypoxia imaging was accomplished for a 6×6 cm field of view (FOV). The FOV was illuminated by a 635nm pulsed LED (SOLIS-620D, Thorlabs). To provide higher wavelength specificity the illumination was coupled to a 635nm band-pass filter. The illumination used a pulse-width of 225 microseconds at a frequency of 2kHz with power density of 5mW/cm2. A 635nm illumination is chosen as lower absorption and scattering from tissue chromophores are observed in this band.^51^ This would allow for penetration depth greater than 0.5 mm, which is the approximate thickness of mouse skin.^51^ To collect DF hypoxia signals, the setup employed a time-gated emICCD camera (PIMAX-4, Teledyne Princeton Instruments). The camera was synchronized to illumination pulses and was time-gated to capture after each illumination pulse to avoid PpIX PF and collect only PpIX DF. The camera collected at a gate-width of 250 microseconds as previous work has shown how temporal oversampling aids to increase the collected DF signal intensity.^52^ As the DF signal is amplified visualizing features with less hypoxia/DF intensity becomes plausible. To isolate collected PpIX fluorescence a 698/70 filter (FF01-698/70-25, Semrock) was used in the detection path. This allowed for the second major peak of PpIX to be collected in a wavelength range with low tissue chromophore absorption and scattering. This is referred as the hypoxia or DF channel throughout this manuscript. A dichroic mirror was placed in the path of PpIX detection together with CMOS sensors 1 and 2 (Blackfly, FLIR, Teledyne Technologies) to concurrently capture both white light and PF signals synchronized to the time-stamps of the DF channel. Videos were collected at an effective frame rate of 20 fps, allowing for real-time imaging of hypoxia. Image frames were acquired with a pixel resolution of 1024 by 1024 pixels and a 2×2 binning applied before image analysis. In summary the setup allows for concurrent imaging of white-light (RGB), PpIX PF and DF at real-time frame rates. These data sets were saved as MP4 and AVI video files containing all frames recorded.

### C. MICE ANESTHESIA PROCEDURES

In vivo experiments were used to validate the biology using Athymic nude mice (Envigo/Inotiv, Inc) in vivo in five different scenarios: Moderate wounds, visually inflamed wounds, mice with no wounds, mice in movement and mice bearing AsPC1 Pancreatic Adenocarcinoma tumor xenografts. With the exception of mice in movement, mice were anesthesized (SomnoSuite Low-Flow anesthesia system, Kent Scientific) for wound procedures, 5-ALA injection and for imaging sessions. Anesthesia was done with a flow rate of 3% isofluorane for induction and 2-2.5% for maintainance, using 100% oxygen (500 ml/min) as the carrier gas. (SomnoSuite Low-Flow anesthesia system, Kent Scientific)

### D. DEVELOPMENT OF WOUND MODELS IN MICE

To develop both moderate (N=6) and visually inflammed wound models (N=6), mice were anesthesized and wounds inflicted on the flank area of the right leg. For moderate wounds, surgical scissors were used to cut a wound of *∼* 5 mm length and approximate depth of 3 mm. More severe wounds included an incision of 5 mm length and estimated depth of 5 mm. Wounds were subsequently sutured and monitored over the mentioned time-points. For analgesia after wounds are inflicted, Ketoprofen is administered at a dose of 20 mg/kg through subcutaneous injection. After wounds and anesthesia recovery, mice are monitored in the recovery cage for any signs of distress. For imaging time-points visually inflammed mice are classified into this category, while non-inflamed mice are classified into moderate wounds. A control group of mice with no wounds is also imaged. The wounded mice were housed at UW-Madison vivarium, according to UW Madison Animal Protocol Number M006554-R01-A01 and received a chlorophyll-free purified diet (TD.97184, Envigo RMS LLC) to reduce autofluorescence from the skin surface, colon and feces.

### E. DEVELOPMENT OF ASPC1 TUMOR XENOGRAFTS

To develop AsPC1 pancreatic adenocarcinoma tumor xenografts in mice, the same protocol previously reported was used. ^34^ Cells acquired from ATCC were cultured in RPMI 1640 media supplemented with 1% penicillin/streptomycin and 10% fetal bovine serum. To grow cells a 5% CO2 incubator at 37°C was used. The cells were collected and resuspended in a mixture of 50% Matrigel / 50% PBS. Cells were injected into the flank of the right leg of athymic nude mice (6-8 weeks of age, Envigo) at a concentration of 1×106 cells / 0.1mL injection. Tumor size was monitored and when a 7-8mm diameter was reached (*∼* 4 weeks after) mice were imaged. Under anesthesia, mice (N=6) received the weight based 5-ALA injection 1 hour prior to imaging. The tumor bearing mice were housed at UW-Madison vivarium, according to UW Madison Animal Protocol Number M006554-R01-A01 and received received a chlorophyll-free purified diet (TD.97184, Envigo RMS LLC) to reduce autofluorescence from the skin surface, colon and feces.

### F. PPIX DF WOUND IMAGING AND COMPARISON TO ICG BASED LOCALIZATION

5-ALA was injected retro-orbitally at 250 mg/kg dose for all groups of mice. This dose was calculated as a human-equivalent dose using a mg/m^2^ conversion from humans to mice, resulting in the mouse dose that is equivalent to 20 mg/kg in a human. ^53^ PpIX imaging was then performed at 1, 3 and 6 hours post 5-ALA administration for a visually inflamed wounded mice group (N=4). This was done to understand time-points at which DF intensity signal was the highest. After determination of highest DF intensity timepoint, PpIX DF imaging was conducted under anesthesia at 1 hr post 5-ALA injection in all cases. Groups of wounded mice (N = 6) were then imaged at day 1, 3, 5 and 7 post intial wound. This was done with the goal to assess differences in lymphatic response across multiple days post-wound. Control mice with no wounds (N = 6) were also imaged and used as experimental controls and 2 mice from each group also imaged under movement. Each mouse was dynamically imaged for a span of 10-15 minutes. The location of pulsating node regions imaged through DF lymphatic imaging, specifically SLN and PLN nodes was correlated to location areas from ICG. After non-invasive PpIX imaging of the mice through the skin, and while the mouse was under anesthesia, indocyanine Green (ICG) at a concentration of 0.1 mg/mL was injected subcutaneously into the second dorsal toe of the hindfoot, with the needle oriented rostrally.^22^ The ICG contrast was used to non-invasively to confirm the location of the lymphatic vessels and nodes proximal to the right hind-paw.^20^ The injection site was expected to form a small bleb before ICG was gradually absorbed into the lymphatic vessels. This process was expedited by gently massaging the mouse’s hind-paw. After ICG injection, the lymph nodes under the skin were exposed by carefully dissecting and removing the skin from the center of the back of the mouse to the dorsal region of the mouse (Figure 1 (g)). ICG was imaged and subsequently PpIX DF was also imaged for the dissected mouse. ICG imaging was performed through the EleVision IR with Visionsense VS3 Iridium platform (Medtronic, Minneapolis, MN). The center of the FOV for the ICG imager was aligned to match the center of the FOV for the PpIX DF imaging system.

### G. CORRELATION OF PPIX DF SIGNALS TO RESPIRATION AND HEART-BEAT RATES

Besides verifying localization of pulsating nodes through ICG, pulsation patterns recorded through DF PpIX imaging were compared to patterns resulting from respiration rate and heart rate. During DF imaging of wounded mice (N=6) they were placed over an electrode based vital sign monitor for rodents (Rodent Surgical Monitor+, Indus Instruments), contact gel was placed between the four paws and electrodes and respiration and heart (Lead I, II, II) rate measured. Pad temperature was set to 35 degrees Celcius. The instrument resolved the real-time heart rate and respiration rate values. Values were averaged over a 10 minute period to match the the DF Imaging time. Start of vital sign monitoring was timed to match start of DF imaging. Averaged values in BPM for heart rate and BrPM for respiration rate were compared to BPM values estimated through the DF imaging measurements. For selected cases and for visualization purposes, waveforms of respiration and ECG were compared to waveforms observed from PpIX DF imaging.

### H. DF IMAGE PROCESSING AND QUANTIFICATION

The data files acquired through the DF Imaging system included MP4 and AVI videos recorded for the duration of measurements. Image processing was performed in MATLAB. Processing included subtraction of background per frame, followed by 2×2 binning of the 1024×1024 pixel resolution, yielding frames with 512×512 resolution for quantification. Additionally, to reduce computational burden on the pulse/beat per minute (BPM) estimation and improve image quality, frames were averaged every 5 frames. A region selector GUI was developed to freely select areas of interest. For quantification of BPM, the DF imaging dataset, which contained lymphatic activity information, was used.

In cases of correlation to ICG since the center of the FOV is aligned but not the FOV size, image registration is performed in relation to the silhouette of the mouse and location of wound. For wound sets, regions in the wound and in some cases inflammation regions, as well as SLN and PLN node regions were selected and average temporal sequence extracted over the duration of the video. Zero values were set to NaN before average of the accounted timeframe.

Figure 3 (d) displays an example sequence for one minute. The peaks of the sequence were then calculated per minute and average BPM value used. For cases where mice are in movement since fluctuations can be compromised by movement, quantification was limited to frames where the mouse position was kept approximately constant and to intensity only values. For these frames intensity values were averaged for a window of 20 frames for the SLN and PLN regions and the process repeated for control mice under anesthesia.

For AsPC1 tumor DF datasets, regions of interest were selected at the tumor, SLN and PLN regions and BPM calculated as previously mentioned. For DF intensity estimation frames over the duration of the dataset were averaged and average intensity value calculated for the regions of interest (ROIs) per mouse. The group statistics were plotted as shown in Figure 5(d) for the groups for both SLN, PLN nodes and tumor regions. To understand the relationship between PF and DF time-points, average PF intensity is retrieved for the ROIs as well as the skin area of the mouse and PF tumor to background ratio (TBR) calculated.

## ACKNOWLEDGMENT

The research presented in this article was funded by the NIH grants P01 CA084203 and R01 EB032337.

## AUTHOR CONTRIBUTIONS

M.O. and B.W.P. designed research; M.O., and M.R. performed experiments; M.O., and B.W.P. developed the analytic tools; M.O., SOP, WZ and B.W.P. analyzed the data; M.O. and B.W.P. wrote the paper, and all authors reviewed and revised the manuscript.

