## Supplementary Material for "Lymphatic dynamics visualized by native transient hypoxia imaging in areas of tissue damage, edema and in sentinel lymph nodes"

**Supplementary Text**

The pre-clinical DF hypoxia imaging setup is depicted in Supplementary Figure 1 (S1). Macroscopic imaging of PpIX DF hypoxia imaging was accomplished for a 6 x 6 cm field of view (FOV). The FOV was illuminated by a 635nm pulsed LED (SOLIS-620D, Thorlabs). To provide higher wavelength specificity the illumination was coupled to a 635nm band-pass filter. The illumination used a pulse-width of 225 microseconds at a frequency of 2kHz with power density of 5mW/cm2. To collect DF hypoxia signals, the setup employed a time-gated emICCD camera (PI-MAX-4, Teledyne Princeton Instruments).

The camera was synchronized to illumination pulses and was time-gated to capture after each illumination pulse to avoid PpIX PF and collect only PpIX DF. The camera collected at a gate-width of 250 microseconds as previous work has shown how temporal oversampling aids to increase the collected DF signal intensity.[51] To isolate collected PpIX fluorescence a 698/70 filter (FF01-698/70-25, Semrock) was used in the detection path. This allows for the second major peak of PpIX to be collected in a wavelength range with low tissue chromophore absorption and scattering. This is referred to as the hypoxia or DF channel throughout this manuscript. A dichroic mirror is placed on the path of PpIX detection together with CMOS sensors 1 and 2 (FLIR, Blackfly, Teledyne Technologies) to concurrently capture both whitelight and PF signals synchronized to the time-stamps of the DF channel. Videos are collected at an effective frame rate of 20 fps, allowing for real-time imaging of hypoxia. Image frames were acquired with a pixel resolution of 1024 by 1024 pixels and a 2x2 binning applied before image analysis. In summary the setup allows for concurrent imaging of white-light (RGB), PpIX PF and DF at real-time frame rates. These data sets are saved as MP4 and AVI video files containing all frames recorded.

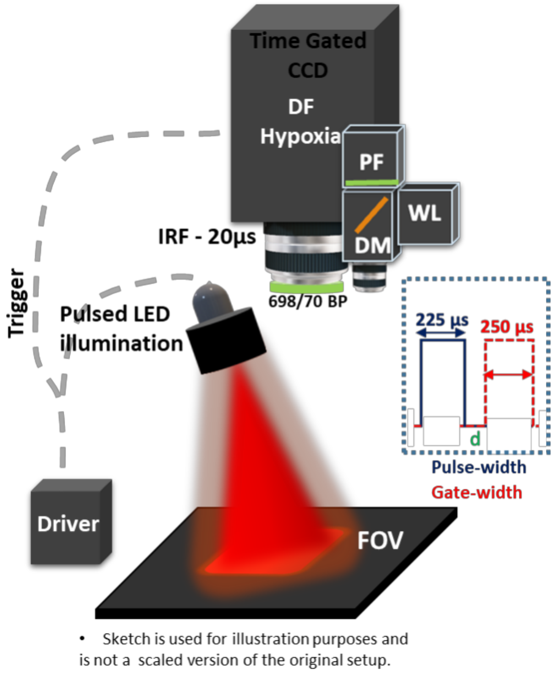


**Fig. S1.** Schematic of system and acquisition settings used for imaging of PpIX DF hypoxia,

Supplementary Figure 2 (S2) describes the DF hypoxia response when the mouse model is restrained in a container vial and skin is in contact with the container surface. Hence for experiments with freely moving mice, the mice were only hold by the tail area and outside of the FOV to avoid induced hypoxia effects on the skin. The response of skin to pressure has been previously reported.

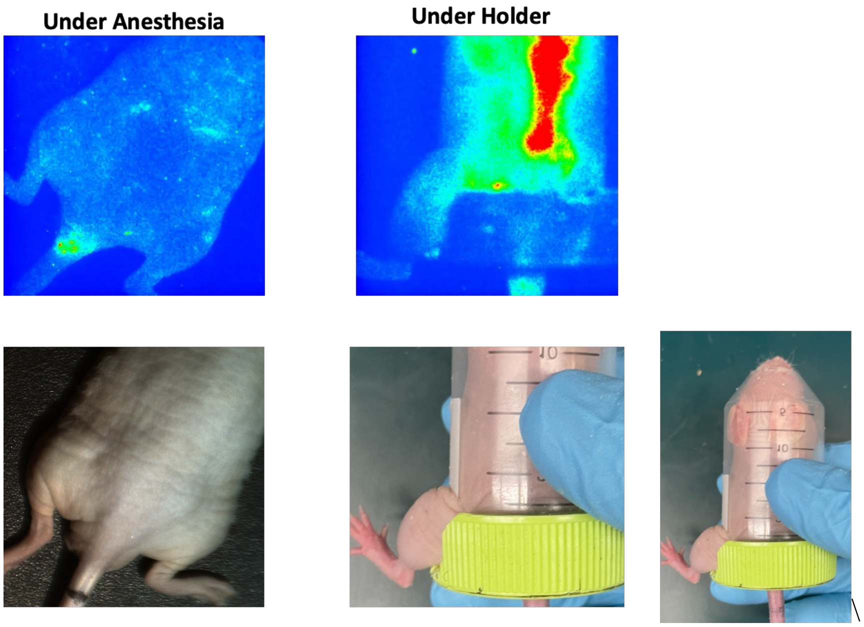


**Fig. S2.** Evidence of skin response when skin is pressured or covered through a mouse holder. DF hypoxia signal and white light images displayed for two scenarios. When the mouse is under anesthesia and when the same mouse is freely moving within a restrainer vial.

**Video S1 (separate file).**

Lymphatic response as recorded by PpIX DF imaging for a mouse with a moderate level wound injury.

**Video S2 (separate file).**

Example of lymphatic response as recorded with PpIX DF imaging for a non-injured mouse with and without anesthesia (awake and moving).

**Video S3 (separate file).**

Example of lymphatic response as recorded with PpIX DF imaging for a severe injured mouse model.

**Video S4 (separate file).**

Example of lymphatic response seen in DF Hypoxia for both injured mouse model across different days post-wound.

**Video S4_2 (separate file).**

Example of lymphatic response seen in DF Hypoxia for injured mouse model with visible inflammation across different hours post-5ALA injection.

**Video S5 (separate file).**

Lymphatic response as recorded with PpIX PF and DF imaging for a mouse with AsPC1 tumor xenograft across different timepoints post 5-ALA injection.
